# An architectonic type principle in the development of laminar patterns of cortico-cortical connections

**DOI:** 10.1101/866681

**Authors:** Sarah F. Beul, Alexandros Goulas, Claus C. Hilgetag

**Author notes:** Shared senior authors (Alexandros Goulas, Claus C. Hilgetag).

## Abstract

Structural connections between cortical areas form an intricate network with a high degree of specificity. Many aspects of this complex network organization in the adult mammalian cortex are captured by an architectonic type principle, which relates structural connections to the architectonic differentiation of brain regions. In particular, the laminar patterns of projection origins are a prominent feature of structural connections that varies in a graded manner with the relative architectonic differentiation of connected areas in the adult brain. Here we show that the architectonic type principle is already apparent for the laminar origins of cortico-cortical projections in the immature cortex of the macaque monkey. We find that prenatal and neonatal laminar patterns correlate with cortical architectonic differentiation, and that the relation of laminar patterns to architectonic differences between connected areas is not substantially altered by the complete loss of visual input. Moreover, we find that the amount of change in laminar patterns that projections undergo during development varies in proportion to the relative architectonic differentiation of the connected areas. Hence, it appears that initial biases in laminar projection patterns become progressively strengthened by later developmental processes. These findings suggest that early neurogenetic processes during the formation of the brain are sufficient to establish the characteristic laminar projection patterns. This conclusion is in line with previously suggested mechanistic explanations underlying the emergence of the architectonic type principle and provides further constraints for exploring the fundamental factors that shape structural connectivity in the mammalian brain.

## Introduction

Brain regions are linked by a complex, characteristic network of structural connections. One principle that has been shown to reliably capture multiple features of cortico-cortical connectivity in the adult mammalian brain is the structural model (Barbas 1986, 1995; Barbas and Rempel-Clower 1997; García-Cabezas et al. 2019), also called architectonic type principle (Hilgetag et al. 2019), which relates features of mesoscopic structural connections to the relative architectonic differentiation of the brain. The architectonic type principle accounts for data observed in a number of mammalian species, such as the rhesus macaque (Medalla and Barbas 2006; Medalla et al. 2007), the cat (Hilgetag and Grant 2010) or the mouse (Goulas et al. 2017), connections within as well as between the cortical hemispheres (Barbas et al. 2005; Goulas et al. 2017), connections towards the primate frontal cortex (Barbas 1986; Barbas and Rempel-Clower 1997; Rempel-Clower and Barbas 2000; Ghashghaei et al. 2007), within the primate and cat visual system (Hilgetag and Grant 2010; Hilgetag et al. 2016) and connections across the entire cortex of the macaque (Beul et al. 2017; Beul and Hilgetag 2019) and the cat (Beul et al. 2015). It has been suggested that such a widely applicable principle may emerge from spatio-temporal interactions of neural populations during ontogeny, without the essential need of sensory input or other major external influences (Barbas 1986, 2015; Barbas and García-Cabezas 2016; Hilgetag et al. 2016). The experimental study of the formation of structural connections, which happens concurrently with the formation of the brain itself, is an onerous endeavour. Consequently, the first support of the suggested mechanistic explanation for the emergence of the architectonic type principle has come from *in silico* models, where simulation results have shown that simple interactions between the time and place of neurogenesis can result in structural networks that capture many of the relationships observed in empirical mammalian cortico-cortical connections (Beul et al. 2018; Goulas, Betzel, et al. 2019). At present, these simulation experiments have explored how the existence of connections (i.e., whether brain areas are connected or not) and the strength of projections can be shaped. Naturally, further features of connectivity, beyond the basic existence of connections, are of interest. One of these features is presented by the laminar patterns of projection origins, which are strikingly regular (Rockland and Pandya 1979) and well captured by the architectonic type principle (Barbas 1986; Barbas and Rempel-Clower 1997; Rempel-Clower and Barbas 2000). They represent a connectional feature that is fundamental to advanced theories of relations between brain structure and function (Friston 2010; Feldman Barrett and Simmons 2015), but how these laminar patterns are shaped during ontogeny is presently not fully clear. Since the architectonic type principle is the most central predictor of laminar projection patterns in multiple species as documented so far (cf. Barbas 2015; Hilgetag et al. 2019), there are two prominent questions about the origin of this relationship between architectonic differentiation and laminar patterns. First, it is not clear if the relation of architectonic differentiation and laminar origin of connections applies only to the adult state of the cerebral cortex. It was shown that the laminar origin of connections is not uniform, but already biased across areas early in development (Barone et al. 1995). Hence, one may wonder if the architectonic type principle reflects graded differences of the laminar origin of connections already in prenatal and neonatal states of the connectivity, or if the early laminar origin patterns of areas undergo drastic reconfigurations which alter the initial bias and thereby eventually give rise to the architectonic type principle in the adult animal. In addition to the extent to which a biased distribution of laminar origins constitutes a pre-configuration of the adult state, a second question concerns the mechanisms that effect the refinement of laminar projection patterns. These could be intrinsic factors, such as apoptosis, or extrinsic factors, such as synaptic activity resulting from sensory input. Enucleation experiments allow inferring the influence of the visual input on the formation of cortical areas and connections, e.g., Karlen and Krubitzer (2009), and are thus helpful in deciphering the influence of external stimuli on the formation of connectional features.

Making use of tract-tracing data detailing laminar patterns of projection origins obtained in the immature macaque cortex (Kennedy et al. 1989; Batardière et al. 2002; Magrou et al. 2018), we here investigate the extent to which the architectonic type principle applies to connectional data from early development and enucleated animals, presenting findings which indicate that processes very early during ontogenesis are sufficient to establish laminar projection patterns that are consistent with the architectonic type principle.

## Methods and Results

To assess whether laminar patterns of projection origins were correlated with relative architectonic differentiation of connected areas in the immature cortex of the macaque monkey, we combined five different resources providing measures of laminar projection patterns and architectonic differentiation.

### Projection data

Measures of laminar projection patterns in the developing and adult macaque cortex were taken from previously published reports (Kennedy et al. 1989; Batardière et al. 2002; Chaudhuri et al. 2015; Magrou et al. 2018). Briefly, Kennedy and colleagues (1989) injected retrograde tracers in the neonate and adult striate cortex (area V1) of cynomolgus monkeys (*Macaca irus*). They evaluated labelled projection neurons in the posterior bank of the lunate sulcus (area V2), on the prelunate gyrus (area V4), and in the posterior bank and fundus of the superior temporal sulcus (STS, which we interpreted to correspond to areas FST, PGa and STPi in the M132 parcellation of Markov et al. (2014)). For each observed projection, they determined the fraction of labelled neurons that originated in supragranular layers (*N*_SG_%). Batardière and colleagues (2002) followed a similar approach, injecting retrograde tracer in area V4 of macaque monkeys (*Macaca fascicularis*) at different fetal stages (embryonic day 112 to embryonic day 140) and in adult monkeys. They evaluated labelled projection neurons across 10 brain areas and also determined the proportional contribution of supragranular neurons (*N*_SG_%) to each projection (Batardière et al. (2002), their Figure 7A).

Magrou and colleagues (2018) performed bilateral enucleation (removal of the eyes) in macaque monkey (*Macaca fascicularis*) fetuses between embryonic days 58 and 73. Retrograde tracers were injected into areas V2 and V4 postnatally, at postnatal day 16 and postnatal month 10, respectively. Labelled projection neurons were evaluated across 18 and 16 brain areas, respectively, and the fraction of labelled projection neurons located in supragranular layers (*N*_SG_%) was determined. We compared the contribution from supragranular neurons in enucleated monkeys to *N*_SG_%-values from intact adult macaque monkeys reported by Chaudhuri and colleagues (2015).

All *N*_SG_%-values that we considered in our analyses are summarized in Online Resource 1.

### Measures of architectonic differentiation

We considered two measures of architectonic differentiation, specifically architectonic type and neuron density. Architectonic type is an ordinal measure of differentiation assigned by experts based on a cortical area’s overall appearance in different types of tissue stains, while neuron density is measured stereologically and has been shown to be a very distinctive marker of individual cortical areas (Dombrowski et al. 2001; Beul and Hilgetag 2019). Both measures have been published previously for the cortical areas considered here (Hilgetag et al. 2016) and are strongly correlated with each other (in this sample of areas, Spearman rank-correlation coefficient ρ = 0.96, p = 3.9e-8). Architectonic type was available for all considered areas, and neuron density was available for all 4 areas considered by Kennedy and colleagues (1989), for 10 of the 11 areas considered by Batardière and colleagues (2002), as well as for 14 of the 20 areas considered by Magrou and colleagues (2018). We report results for both measures to present a comprehensive set of observations that is more robust against possible shortcomings of a particular measure.

### Immature projection patterns correlate with adult differentiation measures

When immature (i.e., prenatal and neonatal) laminar patterns of projection origins are compared to their eventual adult composition, a clear correspondence can be observed, such that the bias in origin layers existent in the immature cortex largely persists in the adult cortex (Figure 2A, Online Resource 2). Consistent with this observation, immature patterns of laminar origins are strongly correlated with the difference in architectonic differentiation between connected areas (Figure 2B, C, Online Resource 2). For comparison, we also show the relation between adult *N*_SG_%-values and difference in architectonic differentiation in these panels. Note that the slope of the regression lines becomes steeper for adult laminar patterns compared to immature patterns, indicating that an initial asymmetry in laminar contributions sharpens with maturation. The relation between immature and adult *N*_SG_%-values becomes even clearer in Figure 1D and E (also see Online Resource 2), which show that the amount of remodelling that a projection undergoes from the immature to the adult state is also correlated with the connected areas’ relative architectonic differentiation. This implies that later processes serve to refine a projection’s laminar origins further towards a laminar bias that was already present from the outset.

**Figure 1.**
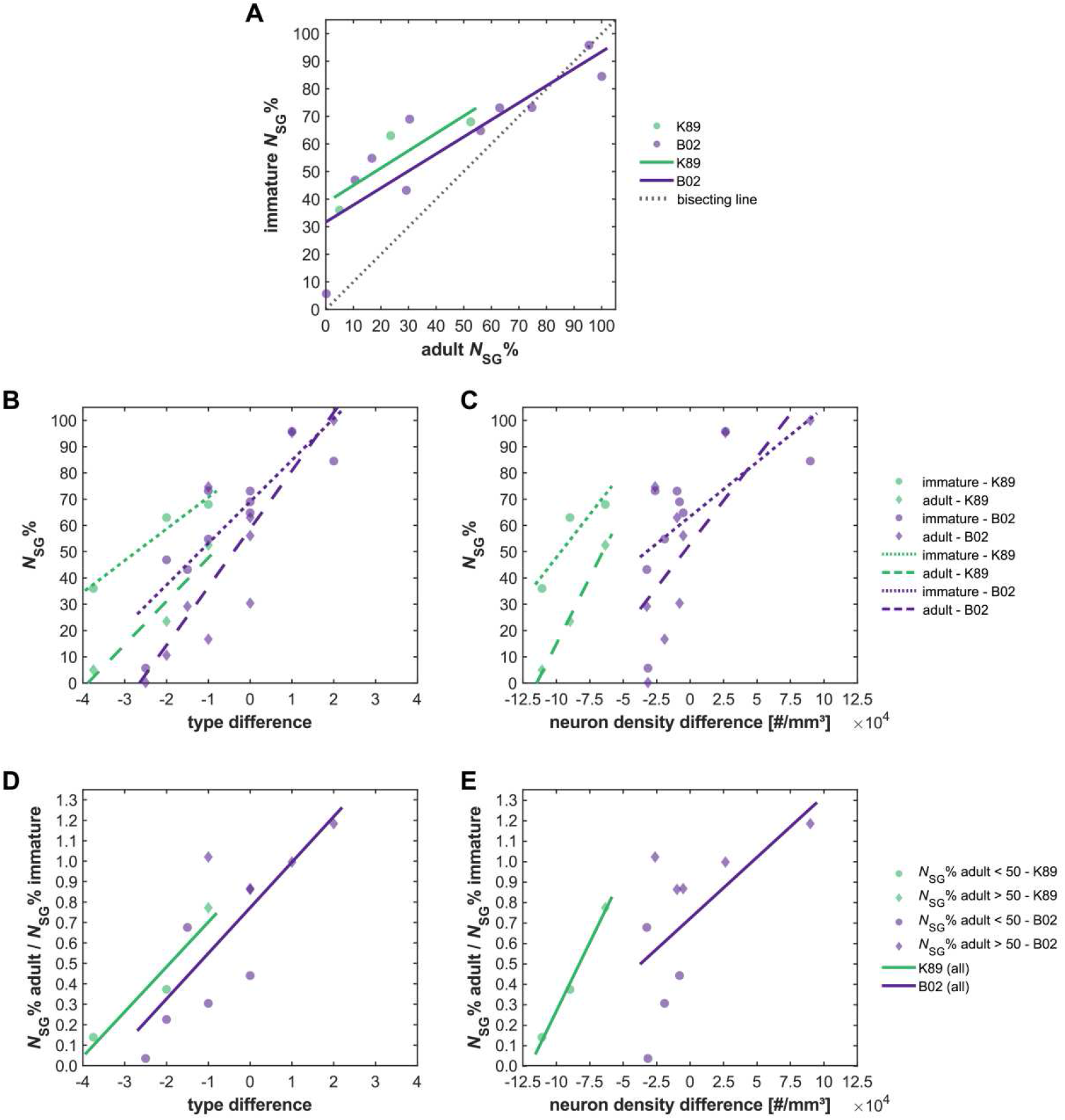
Laminar projection patterns in immature cortex. Relative contribution of supragranular projection neurons (*N*_SG_%) to projections targeting areas V1 (K89, neonatal) and V4 (B02, fetal) in the immature macaque cortex. (A) Immature *N*_SG_% in relation to the corresponding adult *N*_SG_%. (B) *N*_SG_% for both immature and adult cortex in relation to architectonic differentiation measured as difference in architectonic type, where type difference = type_source area_ – type_target area_. (C) *N*_SG_% for both immature and adult cortex in relation to architectonic differentiation measured as difference in neuron density, where neuron density difference = density_source area_ – density_target area_. (D) Fraction of supragranular projection neurons observed in the immature cortex that remains in the adult cortex in relation to difference in architectonic type. (E) Fraction of supragranular projection neurons observed in the immature cortex that remains in the adult cortex in relation to difference in neuron density. Generally, the supragranular contribution declines with maturation. That is, in (D) and (E), the value of adult *N*_SG_% divided by immature *N*_SG_% is below 1 for most areas. Projection data from K89 (Kennedy et al. 1989) and B02 (Batardière et al. 2002).

### Loss of visual input does not substantially alter the gradient of projection patterns

The supragranular contribution to projections in enucleated infant monkeys is strongly correlated with the respective supragranular contribution in intact adult monkeys, especially if connections from highly affected area V1 (cf. detailed descriptions in Magrou et al. (2018))are excluded (Figure 2A, Online Resource 3). However, there is a tendency towards higher supragranular contributions in the enucleated infants (as most data points are above the bisecting line), especially for projections to area V2. Indeed, a permutation test shows the median change in *N*_SG_% (i.e., enucleated *N*_SG_% - intact *N*_SG_%) to be larger for injections in V2 than in V4 (p = 0.01, 10^4^ permutations). Since the tracer was injected at different ages for projections to V2 and V4, the higher supragranular contribution could be explained by differences in maturation: V2 was injected earlier (at postnatal day 16) than V4 (at postnatal month 10), which may have caused the *N*_SG_% values of projections targeting V4 to be more similar to the intact adult *N*_SG_% values. This hypothesis is in line with the generally higher *N*_SG_% values observed for the prenatal and neonate injections reported by Batardière and colleagues (2002) and Kennedy and colleagues (1989). In principle, a comparison with neonatal projection patterns in intact monkeys would have been preferable to a comparison to adult patterns, but these data are not available for the projections between areas reported by Magrou and colleagues (2018). As it is, it might be argued that the projection patterns after enucleation are even less affected than it appears here, since the laminar patterns would likely undergo further postnatal changes, similar to the change already observed for injections in the neonatal to the infant cortex. Extrapolating from our analyses in the previous section, a general decline in supragranular contribution with maturation appears realistic, which would increase the correspondence between the *N*_SG_%-values of intact adults and enucleated infants once they matured to adults.

**Figure 2.**
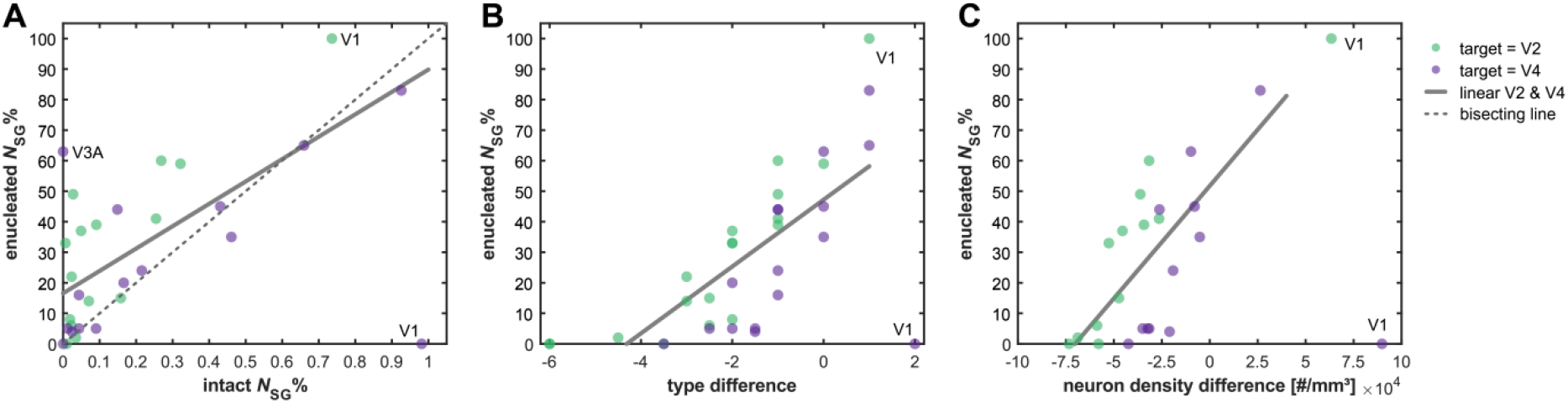
Laminar projection patterns after enucleation. Relative contribution of supragranular projection neurons (*N*_SG_%) to projections targeting areas V2 and V4 in the cortex of enucleated macaque monkeys. (A) *N*_SG_% after enucleation in relation to the respective *N*_SG_% in intact monkeys. (B) *N*_SG_% after enucleation in relation to architectonic differentiation measured as difference in architectonic type, where type difference = type_source area_ – type_target area_. (C) *N*_SG_% after enucleation in relation to architectonic differentiation measured as difference in neuron density, where neuron density difference = density_source area_ – density_target area_. Data from V2 and V4 were pooled for correlations and linear regression. Projections originating in V1 were excluded from the linear regression because V1 was affected very strongly by the enucleation and the resulting *N*_SG_%-values are outliers. Projection data from (Chaudhuri et al. 2015; Magrou et al. 2018).

Despite the drastic effects of enucleation on the organization of the primary visual cortex, the gradual changes in laminar projection patterns that have been reported to align with the relative architectonic differentiation of connected areas can also be observed in enucleated infant monkeys (Figure 2B,C, Online Resource 3). The laminar patterns of projections are strongly correlated with the relative architectonic differentiation of two connected areas, both when it is measured as difference in architectonic type and as difference in neuron density. Thus, despite possible changes in projection patterns, the previously observed relation between laminar patterns and relative differentiation still holds even after complete loss of visual input. In line with the drastic changes to the organization of V1 (cf. Magrou et al. 2018), projections from V1 appear to be altered most strongly. While the correlation of supragranular contribution with architectonic type difference or neuron density becomes stronger if V1 data points are excluded, it is strong and significant even if they are included. This observation implies that the establishment of regular laminar projections patterns is largely independent of normal sensory input, with the possible exception of the directly perturbed areas.

## Discussion

Here we report that the characteristic laminar patterns of cortical projection origins, which have been shown to be closely associated with the relative architectonic differentiation of cortical areas, are already correlated with the later, adult architectonic differentiation in immature macaque monkeys. This observation was made consistently in intact fetal and neonate macaque monkeys as well as in enucleated infant macaque monkeys. Hence, it appears that the processes that determine from which layers a projection originates occur early in development, and are relatively robust to severe changes such as the loss of sensory input. Since we find that laminar projection patterns are consistent with the architectonic type principle soon after their establishment, processes that occur later during ontogenesis may play a smaller role in the emergence of the architectonic type principle. This applies, for example, to processes such as pruning, activity-dependent remodelling or selective apoptosis in the different layers across the cortical gradient. These phenomena may serve to further refine the laminar patterns of projection origins, but do not appear to have a crucial role in determining an overall bias towards infra- or supragranular origins.

These observations concerning immature laminar patterns can inform attempts on explaining how the architectonic type principle may arise during development. Since early, robust processes appear to be sufficient for its emergence, later processes might be omitted from a mechanistic explanation of the origin of the principle without losing much explanatory power. That is, a mechanism that takes into account only early processes but disregards later processes should still be able to generate laminar patterns that do not diverge substantially from empirical observations. We previously demonstrated *in silico* that spatio-temporal interactions in a forming cortical sheet can give rise to connectivity that is consistent with the architectonic type principle in terms of the existence of projections between brain areas (Beul et al. 2018). The fact that immature projection patterns are already consistent with the architectonic type principle, as presented here, implies that such spatio-temporal interactions may also be sufficient for generating the typically observed laminar patterns of projection origins. If the underlying neurogenetic processes can be captured adequately, this link could be demonstrated *in silico*.

Of course, even if laminar projection patterns are determined early in brain development, there remain other candidate mechanisms besides emergence from spatio-temporal interactions. For example, genetic specification is very likely to play some role in the establishment of laminar projection patterns. It has been shown that the eventual projection fate is often acquired even prior to neuronal precursor migration (Jensen and Killackey 1984; McConnell 1988; Polleux et al. 2001) and that initial establishment of connectivity is largely independent of synaptic activity (Verhage et al. 2000). Guidance molecules and their receptors are often expressed in a cell-type specific manner, with many guidance molecules having dual actions depending on the type of receptor to which they bind (Castellani and Bolz 1997; Bagnard et al. 1998; Castellani et al. 1998; Kolodkin and Tessier-Lavigne 2011; Seiradake et al. 2016; Morales and Kania 2017; Stoeckli 2017). These combinations of guidance molecules and receptors have been shown to strongly constrain local, intra-areal connectivity (Bolz and Castellani 1997; Castellani and Bolz 1997). The same principle may apply to longer-range, inter-areal connections. The expression of guidance molecules and receptors is mediated by transcription factors, whose spatially and temporally fine-tuned expression gives rise to distinct cell types with diverse morphological and connectional properties and distinct functions. For example, corticofugal projection identity is mediated by the transcription factors encoded by genes such as Fezf2 and Ctip2, for example reviewed in Molyneaux et al. (2007) and Gaspard and Vanderhaeghen (2011). The effect of Feszf2 expression is not only permissive, but also causal, as forced expression of Fezf2 in progenitors destined for upper layers can induce these cells to atypically project to the pons (Chen et al. 2005). Another example of the genetic specification of a broad class of projection neurons are callosally projecting neurons, of which there are both upper and lower layer populations. Expression of different genes such as Satb2, Hspb3 and Lpl appears to generally specify callosal projection neurons (Alcamo et al. 2008; Molyneaux et al. 2009), while there are also genes specific to either upper or lower layer callosal projection neurons (e.g., Dkk3, Nectin-3 or Plexin-D1 (Molyneaux et al. 2009)). Since there is evidence for the genetic specification of anatomical projection patterns at small (intrinsic, intra-areal circuits) and large (e.g., corticofugal versus callosal projections) spatial scales, projections at intermediate spatial scales, such as cortico-cortical inter-areal projections, are not likely to be an exception from this mode of connection organization. For example, it has been shown that while the white matter of the spinal cord is generally permissive for cortical axon growth, innervation of sections of the spinal grey matter is specific and topographically correct (Stanfield and O’Leary 1985; O’Leary and Stanfield 1986; Kuang and Kalil 1994; Kuang et al. 1994).

To sum up, we draw two main conclusions from the results presented here (Figure 3). First, we show that already in the prenatal and neonatal cortex, the laminar patterns of projection origins correlate with the architectonic differentiation observed in the adult cortex, and that these laminar patterns are not substantially altered by complete loss of visual input. Second, it appears that the initially present biases in laminar projections patterns are progressively strengthened by later developmental processes. During this sharpening of laminar specificity, the amount of change that projections undergo in their supragranular contribution varies concurrently with the relative architectonic differentiation of the connected areas (Figure 3C). These findings have implications for the organization of structural connectivity, indicating that early neurogenetic processes are sufficient to establish the typical laminar projection patterns during brain development. We have previously suggested that the architectonic type principle results from spatio-temporal interactions of neuronal populations in the forming brain (Beul et al. 2018). This mechanism is consistent with the determination of laminar patterns through early neurogenetic processes. Hence, our findings on immature laminar patterns of projection origins strengthen the support for this mechanistic explanation of how the architectonic type principle emerges during ontogenesis.

**Figure 3.**
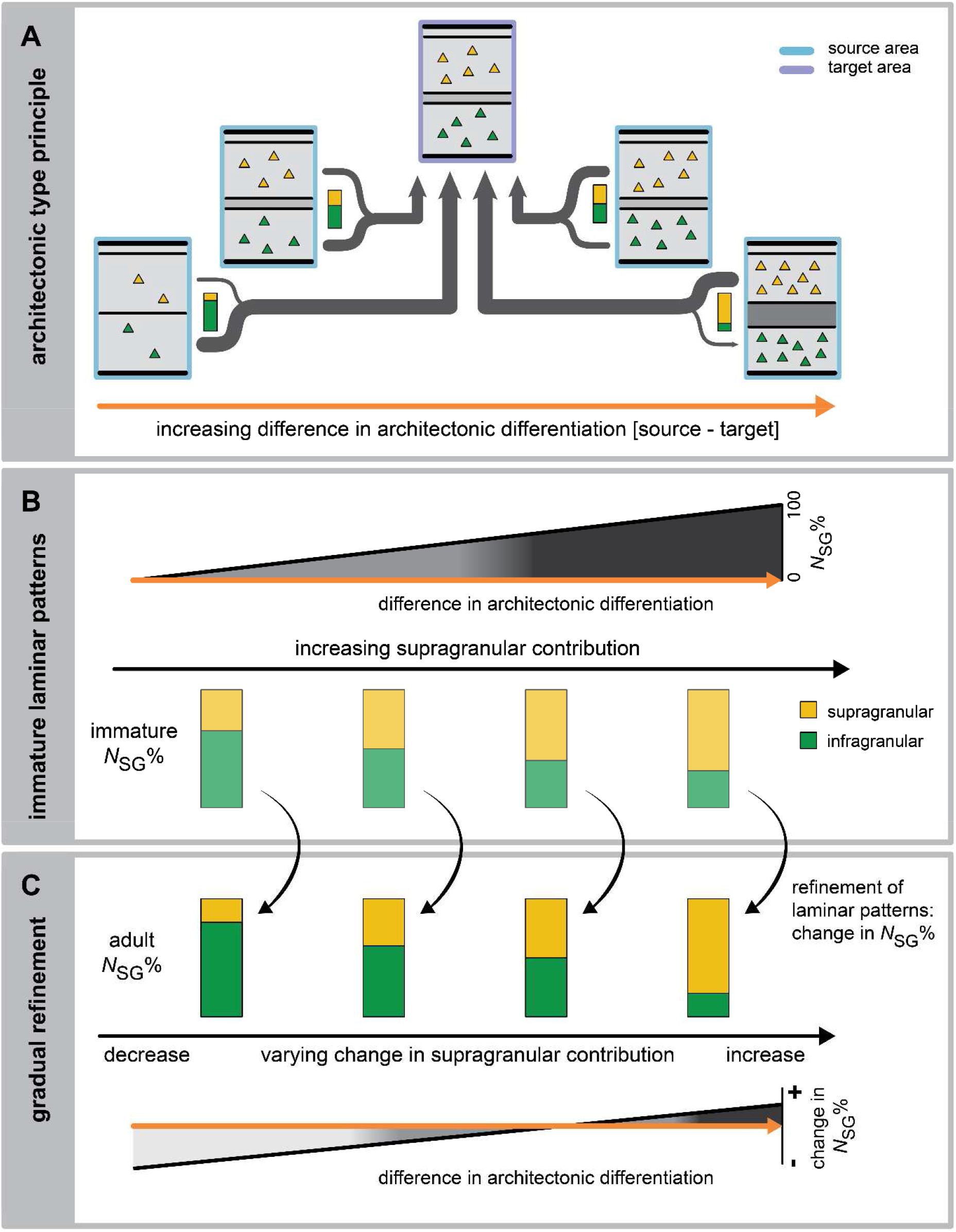
Summary of changes in laminar projection patterns. The architecture of connected cortical areas varies along a natural axis of cortical organization (Hilgetag et al. 2019), such that their relative differentiation may change from a negative to a positive value, as the source area of a projection becomes more differentiated than the target area. (A) The architectonic type principle describes how the proportions of upper and deep laminar origins of projection neurons vary along this gradient of relative differentiation. (B) The present findings show that, already in the immature brain, the contribution of supragranular neurons to a given projection is the stronger, the more differentiated the source area is relative to the target area (immature *N*_SG_%) (cf. Figure 1B, C). (C) This pattern becomes more pronounced as the initially formed projections are refined by later developmental processes (adult *N*_SG_%). Specifically, we observed that this refinement appears proportional to the relative differentiation of connected areas. While the supragranular contribution to projections mainly decreases, the magnitude of this decrease changes concurrently with relative differentiation and eventually reverses into an increase of supragranular contribution. This results in the progressive strengthening of initially present biases in laminar projection patterns (cf. Figure 1D,E).

## Compliance with Ethical Standards

### Funding

This study was supported by the German Research Council (DFG grants SFB 936/ A1, TRR 169/ A2, and SPP 2041 to CCH).

### Ethical approval

This article does not contain any studies with human participants or animals performed by any of the authors.

## Acknowledgements

This work was supported by German Research Council (DFG grants SFB 936/ A1, TRR 169/ A2, and SPP 2041 to CCH).

## Online Resources

**Online Resource 1.**
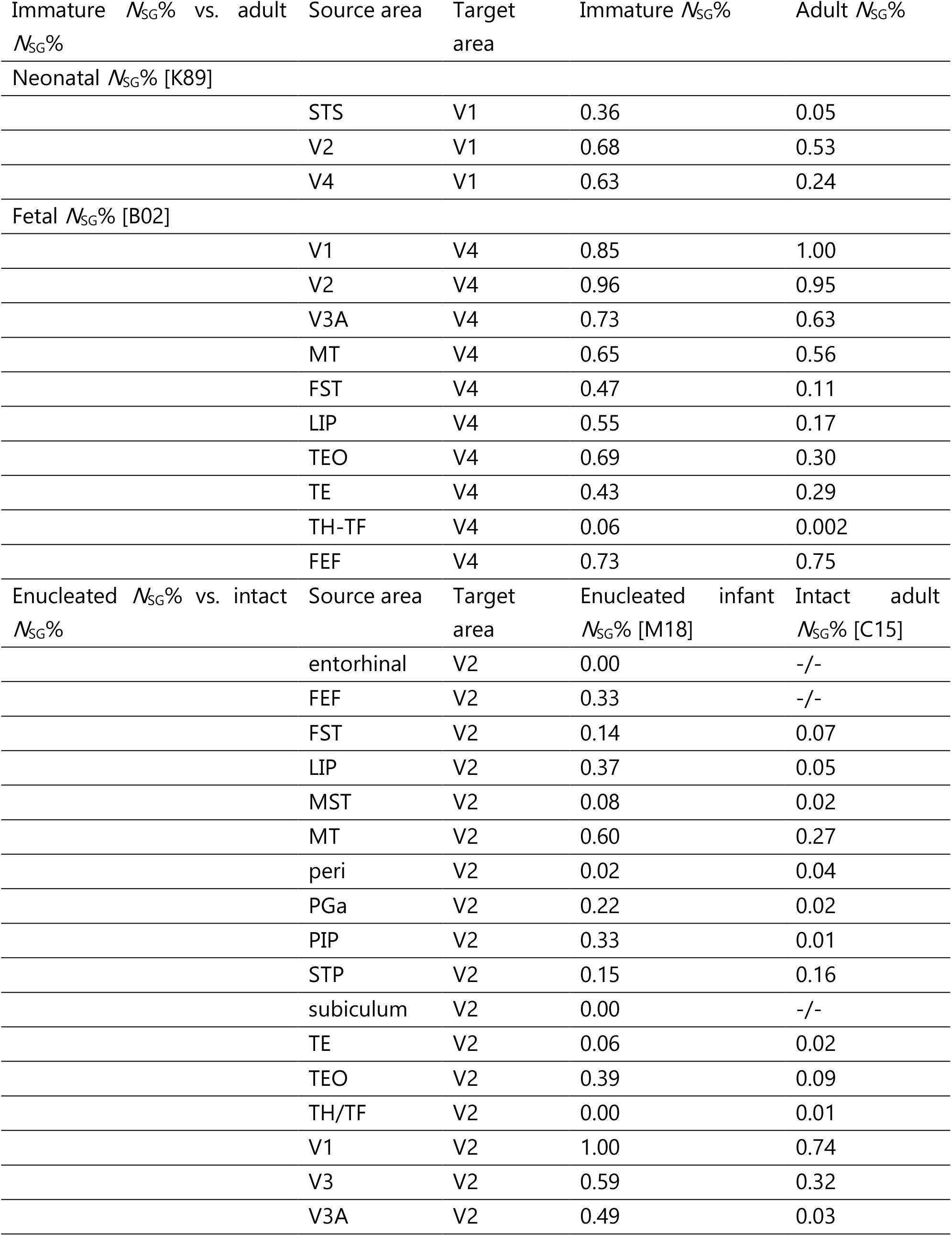
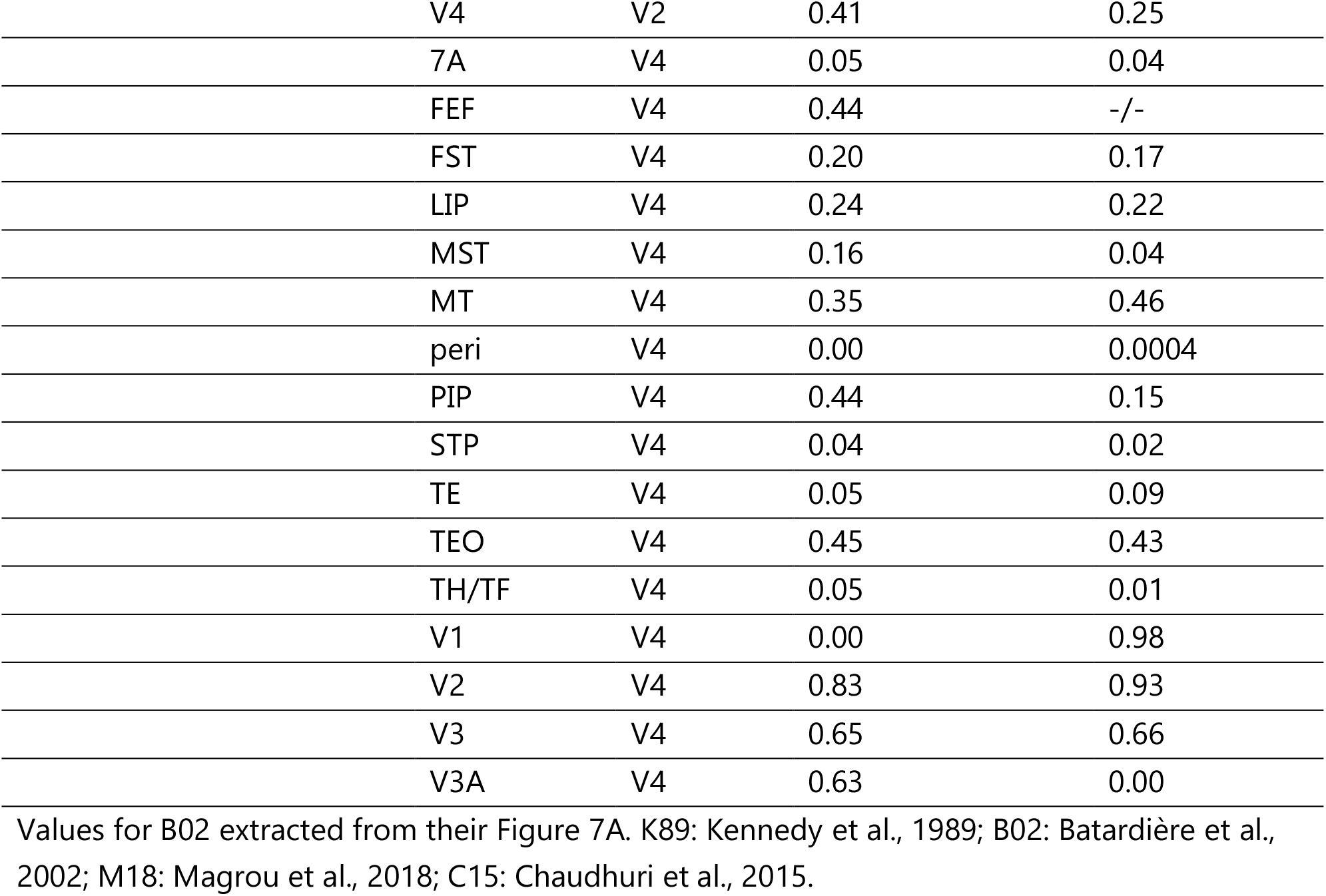
Supragranular contribution to cortico-cortical connections in the immature macaque cortex.

**Online Resource 2.** Correlations of laminar projection patterns and architectonic differentiation in the immature macaque cortex. See Figure 1 for scatter plots of the underlying data. ρ and ρ-value: Spearman rank-correlation; R^2^: coefficient of determination for a linear regression model. Please note that the p-value for K89 correlations cannot be lower than 0.33 because only three data points are available.

| | $\rho$ | p-value | $R^2$ |
| --- | --- | --- | --- |
| adult $N_{SG}\%$ vs. immature $N_{SG}\%$ | | | |
| Neonate (K89) | 1.00 | 0.3333 | 0.77 |
| Fetal (B02) | 0.94 | 0.0000 | 0.75 |
| immature $N_{SG}\%$ vs. architectonic type difference | | | |
| Neonate (K89) | 1.00 | 0.3333 | 0.95 |
| Fetal (B02) | 0.87 | 0.0011 | 0.74 |
| adult $N_{SG}\%$ vs. architectonic type difference | | | |
| Neonate (K89) | 1.00 | 0.3333 | 0.92 |
| Fetal (B02) | 0.87 | 0.0009 | 0.74 |
| immature $N_{SG}\%$ vs. neuron density difference | | | |
| Neonate (K89) | 1.00 | 0.3333 | 0.82 |
| Fetal (B02) | 0.73 | 0.0311 | 0.37 |
| adult $N_{SG}\%$ vs. neuron density difference | | | |
| Neonate (K89) | 1.00 | 0.3333 | 1.00 |
| Fetal (B02) | 0.72 | 0.0369 | 0.54 |
| adult $N_{SG}\%$ / immature $N_{SG}\%$ vs. architectonic type difference | | | |
| Neonate (K89) | 1.00 | 0.3333 | 0.91 |
| Fetal (B02) | 0.77 | 0.0093 | 0.62 |
| adult $N_{SG}\%$ / immature $N_{SG}\%$ vs. neuron density difference | | | |
| Neonate (K89) | 1.00 | 0.3333 | 0.99 |
| Fetal (B02) | 0.58 | 0.1080 | 0.38 |

**Online Resource 3.** Correlations of laminar projection patterns after enucleation. See Figure 2 for scatter plots of the underlying data. ρ and ρ-value: Spearman rank-correlation; R^2^: coefficient of determination for a linear regression model. Projections originating in V1 were excluded because V1 was affected very strongly by the enucleation and the resulting *N*_SG_%-values are outliers.

|  | All data points |  |  | V1 excluded |  |  |
| --- | --- | --- | --- | --- | --- | --- |
| | $\rho$ | p-value | $R^2$ | $\rho$ | p-value | $R^2$ |
| $N_{SG}\%$ enucleated vs. $N_{SG}\%$ intact | 0.47 | 0.0083 | 0.27 | 0.58 | 0.0013 | 0.48 |
| $N_{SG}\%$ enucleated vs. structural type difference | 0.73 | 1.2e-6 | 0.44 | 0.86 | 4.2e-10 | 0.62 |
| $N_{SG}\%$ enucleated vs. neuron density difference | 0.48 | 0.0185 | 0.21 | 0.60 | 0.0033 | 0.47 |

